# Quantitative monitoring of multispecies fish environmental DNA using high-throughput sequencing

**DOI:** 10.1101/113472

**Authors:** Masayuki Ushio, Hiroaki Murakami, Reiji Masuda, Tetsuya Sado, Masaki Miya, Sho Sakurai, Hiroki Yamanaka, Toshifumi Minamoto, Michio Kondoh

**Affiliations:** Department of Environmental Solution Technology, Faculty of Science and Technology, Ryukoku University, Otsu 520-2194, Japan; Joint Research Center for Science and Technology, Ryukoku University, Otsu 520-2194, Japan; Maizuru Fisheries Research Station, Kyoto University, Maizuru, Kyoto 625-0086, Japan; Department of Ecology and Environmental Sciences, Natural History Museum and Institute, Chiba 260-8682, Japan; Graduate School of Science and Technology, Ryukoku University, Otsu 520-2194, Japan; The Research Center for Satoyama Studies, Ryukoku University, Shiga 520-2194, Japan; Graduate School of Human Development and Environment, Kobe University, Hyogo 657-8501, Japan

**Keywords:** biodiversity monitoring, environmental DNA, quantification, internal standard DNA, qPCR, metabarcoding

## Abstract

Effective ecosystem conservation and resource management require quantitative monitoring of biodiversity, including accurate descriptions of species composition and temporal variations of species abundance. Therefore, quantitative monitoring of biodiversity has been performed for many ecosystems, but it is often time- and effort-consuming and costly. Recent studies have shown that environmental DNA (eDNA), which is released to the environment from macro-organisms living in a habitat, contains information about species identity and abundance. Thus, analyzing eDNA would be a promising approach for more efficient biodiversity monitoring. In the present study, we added internal standard DNAs (i.e., known amounts of short DNA fragments from fish species that have never been observed in a sampling area) to eDNA samples, which were collected weekly from a coastal marine ecosystem in Maizuru-Bay, Kyoto, Japan (from April 2015 to March 2016), and performed metabarcoding analysis using Illumina MiSeq to simultaneously identify fish species and quantify fish eDNA copy numbers. A correction equation was obtained for each sample using the relationship between the number of sequence reads and the added amount of the standard DNAs, and this equation was used to estimate the copy numbers from the sequence reads of non-standard fish eDNA. The calculated copy numbers showed significant positive correlation with those determined by quantitative PCR, suggesting that eDNA metabarcoding with standard DNA enabled useful quantification of eDNA. Furthermore, for samples that show a high level of PCR inhibition, our method might allow more accurate quantification than qPCR because the correction equations generated using internal standard DNAs would include the effect of PCR inhibition. A single run of Illumina MiSeq produced > 70 quantitative fish eDNA time series in our study, showing that our method could contribute to more efficient and quantitative monitoring of biodiversity.

## 1. Introduction

Effective ecosystem conservation and resource management require quantitative monitoring of biodiversity, including accurate descriptions of species composition and temporal variations of species abundance. Accordingly, quantitative monitoring of biodiversity has often been performed for many ecosystems. For example, fishing (in aquatic ecosystems), the camera/video trap method (in terrestrial ecosystems) and direct visual census (in aquatic and terrestrial ecosystems) have traditionally been used as tools for biodiversity monitoring [see, for example, 1,2]. These data are invaluable in conservation ecology, but at the same time, the traditional approaches are usually time- and effort-consuming and costly. In addition, most of the traditional methods require professional expertise such as taxonomic identification skill in the field. These difficulties prevent the collection of quantitative, comprehensive (i.e., multispecies and fine-time resolution) and long-term monitoring data about biodiversity.

Environmental DNA (eDNA), which designates DNA isolated from environmental samples (e.g., water or soil) without sampling target (macro-)organism(s), has been used to detect the presence of macro-organisms, particularly those living in an aquatic environment [e.g., 3-5]. In the case of macro-organisms, eDNA originates from various sources such as metabolic waste or damaged tissue [6], and the eDNA contains information about the species identity of organisms that produced it. Since the first application of eDNA analysis to natural ecosystems [7], eDNA in aquatic ecosystems has been used in many studies as a tool for investigation of the distributions of fish species in ponds, rivers and seawater [8–12] as well as the distributions of other aquatic/semiaquatic/terrestrial organisms [13–17]. Recently, researchers have begun to apply high-throughput sequencing technology (e.g., Illumina MiSeq) and universal primer sets to eDNA studies [3–5,14,18]. A previous study demonstrated that an eDNA metabarcoding approach using fish-targeting universal primers (MiFish primers) enabled detection of more than 230 fish species from seawater in a single study [3]. Accordingly, the eDNA metabarcoding approach has become a cost- and labor-effective approach for estimating aquatic biodiversity.

Though the eDNA metabarcoding approach has greatly improved the efficiency of biodiversity monitoring, several potential limitations prevent its use as a tool for quantitative monitoring of biodiversity. First, whether the quantity of eDNA is a reliable index of the abundance (or biomass) of macro-organisms is still controversial. Second, even if the quantity of eDNA is an index of the abundance/biomass of macro-organisms, the number of eDNA sequence reads obtained by high-throughput sequencing may not be an index of the quantity of eDNA, and thus we cannot estimate the quantity of eDNA in an environment by the eDNA metabarcoding approach.

Regarding the first issue, some studies showed that eDNA quantity could be a proxy of the abundance or biomass of macro-organisms under particular conditions such as a tank experiment [e.g., 11]. Also, a recent study showed that eDNA quantity is a proxy of the abundance of fish even in an open ocean ecosystem if appropriate spatial information is incorporated [19]. Thus, there have been many reports of positive and significant linear relationships between eDNA quantity and the abundance/biomass of macro-organisms [11,19–21]. Nonetheless, the general use of eDNA as a proxy of fish abundance/biomass is still controversial because factors that influence eDNA quantity, such as eDNA decay rates in an environment and their release rates from target organisms, are likely to depend on the ecology and physiology of target species and other biotic/abiotic factors [22–24] and because the eDNA quantity found in a sample could also be influenced by water flow in an environment (i.e., eDNA transport). The findings of such studies imply that an accurate estimation of organism abundance/biomass requires sample-specific calibrations that appropriately take into account biotic/abiotic factors (e.g., fish physiological conditions, water temperature, water flow and spatial information). Altogether, the above evidence suggested that information about the abundance/biomass is “encoded” in the quantity of eDNA at least to some extent, and that we may use the quantity of eDNA as “a rough index” of abundance/biomass, but careful interpretations are necessary especially when other related information (e.g., physicochemical properties of water and the ecology of target species) is not available.

Regarding the second issue, some potential approaches to solve this problem have been reported in the field of microbial ecology. For example, Smets et al. [25] added an internal standard DNA (DNA of a microbial species that had never been found in a sample) of known quantity to a soil sample. They used the number of sequence reads of the internal standard DNA to estimate the sequence reads per number of DNA copies, and converted the sequence reads of DNAs from unknown microbial species (i.e., non-standard microbial species) to the number of microbial DNA copies. The total number of microbial DNA copies estimated was significantly positively correlated with other reliable and quantitative indices of soil microbial abundance. Although the quality and quantity of microbial DNA from soil samples could be different from those of macrobial (e.g., fish) eDNA from water samples, this approach is potentially useful to resolve the second issue.

In the present study, we focused on the second issue, i.e., the quantification of eDNA using high-throughput sequencing, and did not explicitly try to resolve the first issue, i.e., the accurate estimation of species abundance/biomass based on eDNA quantity, because at present various biotic/abiotic factors at fine spatiotemporal resolutions, for some of which data are not currently available in the study region, should be incorporated to resolve this first issue [19].

We applied the internal standard DNA method to the eDNA metabarcoding approach to enable quantitative monitoring of multispecies fish eDNA in a coastal marine ecosystem (i.e., identification of fish species and quantification of the number of fish eDNA copies simultaneously). Water samples were collected weekly from a sampling station in Maizuru Bay, located on the Japan Sea coast of central Japan, and eDNAs were extracted from the samples. We added known quantities of short DNA fragments derived from five fish species that have never been observed in the sampling region (freshwater fish species in Southeast Asia or Africa) as internal standard DNAs to each eDNA sample. Using the relationships between the quantity and sequence reads (generated by Illumina MiSeq) of the internal standard DNAs, the sequence reads were converted to the calculated DNA copy numbers. We tested the reliability of our internal standard DNA method by comparing the calculated DNA copy numbers with those quantified by quantitative PCR (qPCR) using several statistical methods. Specifically, we tested the following: 1) whether numbers of sequence reads of the internal standard DNAs linearly correlate with their quantity (copy numbers), 2) whether there is a positive and significant relationship between the calculated DNA copy numbers and those quantified by qPCR, and 3) whether temporal dynamics shown by the internal standard method are comparable with those shown by qPCR.

## 2. Methods

### 2.1. Study site

Water samples were collected at a floating pier in the Maizuru Fishery Research Station of Kyoto University (Nagahama, Maizuru, Kyoto, Japan: 35°28′N, 135°22′E; Fig. 1). The sampling point was located 11 m from the shore, with a bottom depth of 4 m. The adjacent area included a rocky reef, brown algae macrophyte and filamentous epiphyte vegetation, live oysters (*Crassostrea gigas*) and their shells, a sandy or muddy silt bottom and an artificial vertical structure that functioned as a fish reef. The surface water temperature and salinity in the area ranged from 1.2 to 30.8°C and from 4.14 to 34.09‰, respectively. The mean (±SD) surface salinity was 30.0 ± 2.9 ‰ (n = 1,753) and did not show clear seasonality. Further information on the study area is available in Masuda et al. [1, 26].

**Figure 1.**
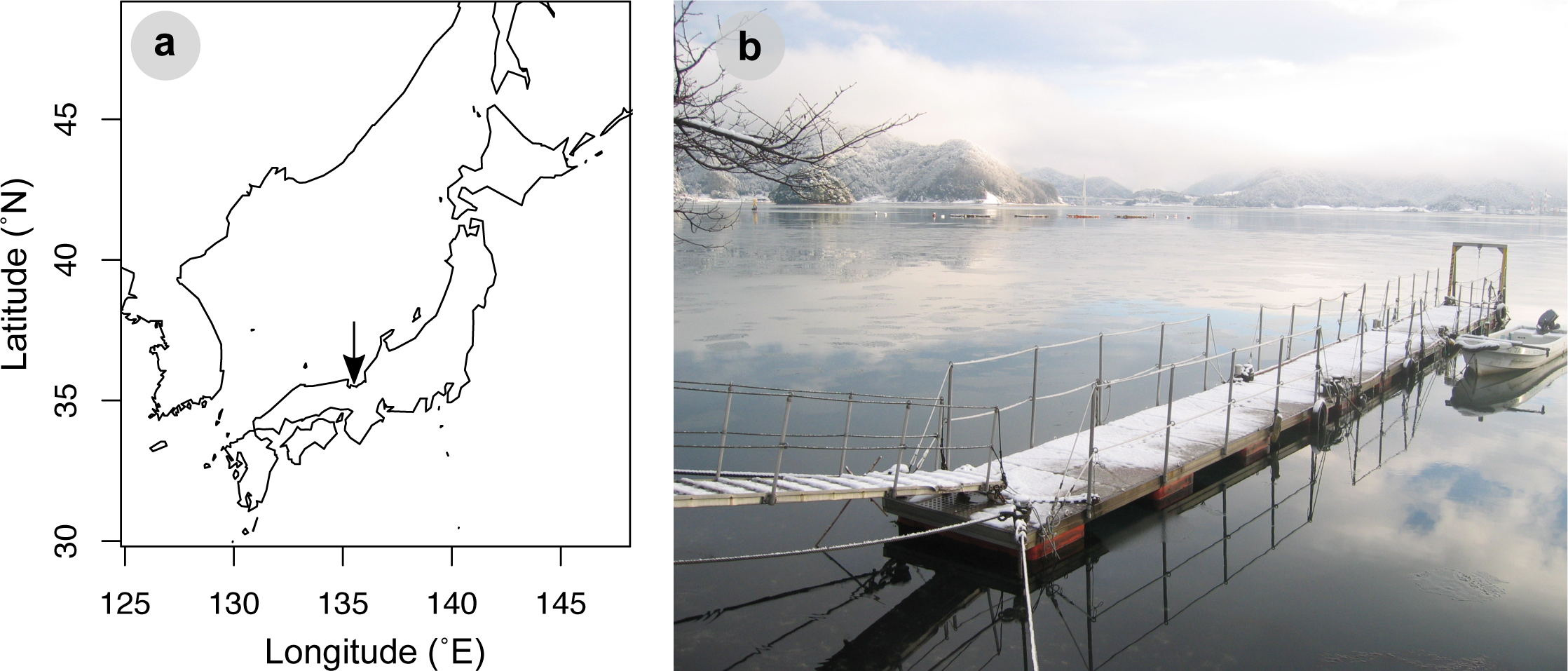
Location of the research site (**a**). The arrow indicates our research site. A floating pier in the Maizuru Fishery Research Station of Kyoto University, Maizuru, Kyoto, Japan, where the weekly water sampling was performed (**b**). Photo taken in winter by R. Masuda.

### 2.2. Water sampling and DNA extraction

All sampling and filtering equipment was washed with a 10% commercial bleach solution before use. We collected 1,000 ml of seawater once a week from 7th April 2015 to 29th March 2016 from a pier (Fig. 1b) in the study area using a polyethylene bottle. Thus, the total number of eDNA samples (excluding artificial seawater samples as negative controls) was 52. The collected water samples were immediately taken to the laboratory and filtered using 47-mm diameter glass-fibre filters (nominal pore size, 0.7 μm; Whatman, Maidstone, UK). The sampling bottles were gently shaken before the filtration. After the filtration, each filter was wrapped in commercially available aluminium foil and stored at –20°C before eDNA extraction. Artificial seawater (1,000 ml) was used as the negative control, and sampling bottles filled with artificial seawater were treated identically to the eDNA samples in order to monitor contamination during the bottle handling, water filtering and subsequent DNA extraction. Negative control samples were obtained once a month (a total of 12 negative controls), and all negative controls produced a negligible number of sequences (i.e., the average number of sequence reads was 2,305 for environmental samples, while it was 21 for negative controls; see Table S1).

DNA was extracted from the filters using a DNeasy Blood and Tissue Kit (Qiagen, Hilden, Germany) in combination with a spin column (EZ-10; Bio Basic, Markham, Ontario, Canada). After removal of the attached membrane from the spin column (EZ-10), the filter was tightly folded into a small cylindrical shape and placed in the spin column. The spin column was centrifuged at 6,000 g for 1 min to remove excess water from the filter. The column was then placed in the same 2-ml tube and subjected to cell lysis using proteinase K. For the lysis, sterilized H_2_O (200 μl), proteinase K (10 μl) and buffer AL (100 μl) were mixed, and the mixed solution was gently pipetted onto the folded filter in the spin column. The column was then placed on a 56°C preheated aluminium heat block and incubated for 30 min. After the incubation, the spin column was centrifuged at 6,000 g for 1 min to collect DNA. In order to increase the yield of DNA from the filter, 200 μl of sterilized TE buffer was gently pipetted onto the folded filter and the spin column was again centrifuged at 6,000 g for 1 min. The collected DNA solution (about 500 μl was purified using a DNeasy Blood and Tissue Kit following the manufacturer’s protocol. After the purification steps, DNA was eluted with the elution buffer (100 μl provided in the kit.

### 2.3. Preparation of standard fish DNAs

Extracted DNAs of five fish species (*Saurogobio immaculatus, Elopichthys bambusa, Carassioides acuminatus, Labeo coubie*, and *Acanthopsoides gracilentus*) that are all freshwater fishes from Southeast Asia or Africa and have never occurred in the sampling region were used as internal standard DNAs. A target region (mitochondrial 12S rRNA) of the extracted DNA was amplified using MiFish primers (without MiSeq adaptors), and the amplified and purified target DNA (about 220 bp) was excised using E-Gel SizeSelect (ThermoFisher Scientific, Waltham, MA, USA). The DNA size distribution of the library was estimated using an Agilent 2100 BioAnalyzer (Agilent, Santa Clara, CA, USA), and the concentration of double-stranded DNA of the library was quantified using a Qubit dsDNA HS assay kit and a Qubit fluorometer (ThermoFisher Scientific, Waltham, MA, USA). Based on the quantification values obtained using the Qubit fluorometer, we adjusted the copy number of the standard DNAs and mixed these DNAs as follows: *S. immaculatus* (500 copiesμl), *E. bambusa* (250 copiesμl), *C. acuminatus* (100 copiesμl), *L. coubie* (50 copiesμl), and *A. gracilentus* (25 copiesμl). Hereafter, the mixed standard DNA is referred to as ‘standard DNA mix’. The numbers of internal standard DNA copies added to samples were determined by quantification of the number of total fish eDNA copies (i.e., MiFish primer target region) using the SYBR-GREEN quantitative PCR method (see below for the detailed method).

### 2.4. Paired-end library preparation

Work-spaces and equipment were sterilized prior to the library preparation, filtered pipet tips were used, and separate rooms were used for pre- and post-PCR operations to safeguard against cross-contamination. We also employed negative controls to monitor contamination during the experiments.

The first-round PCR (1st PCR) was carried out with a 12-μl reaction volume containing 6.0 μl of 2 × KAPA HiFi HotStart ReadyMix (KAPA Biosystems, Wilmington, WA, USA), 0. 7 μl of each primer (5 μM), 0.6 μl of sterilized distilled H_2_O, 2 μl of standard DNA mix and 2.0 μl of template. Note that we included the standard DNA mix for each sample. The final concentration of each primer (MiFish-U-F/R) was 0.3 μ?. The sequences of MiFish primers are: GTC GGT AAA ACT CGT GCC AGC (MiFish-U-F) and CAT AGT GGG GTA TCT AAT CCC AGT TTG (MiFish-U-R). MiSeq sequencing primers and six random bases (N) were combined with MiFish-U primers [see 14 for detailed sequences]. The six random bases were used to enhance cluster separation on the flowcells during initial base call calibrations on the MiSeq platform. The thermal cycle profile after an initial 3 min denaturation at 95°C was as follows (35 cycles): denaturation at 98°C for 20 s; annealing at 65°C for 15 s; and extension at 72°C for 15 s, with a final extension at the same temperature for 5 min. We performed triplicate 1st PCR, and the replicates were pooled in order to mitigate the PCR dropouts. Each pooled 1st PCR product (i.e., one pooled 1st PCR product per sample) was purified using Exo-SAPIT (Affymetrix, Santa Clara, CA, USA). The pooled, purified, and 10-fold diluted 1st PCR products were used as templates for the second-round PCR.

The second-round PCR (2nd PCR) was carried out with a 24-μl reaction volume containing 12 μl of 2 × KAPA HiFi HotStart ReadyMix, 1.4 μl of each primer (5 μM), 7.2 μl of sterilized distilled H_2_O and 2.0 μl of template. Different combinations of forward and reverse indices were used for different templates (samples) for massively parallel sequencing with MiSeq. The thermal cycle profile after an initial 3 min denaturation at 95°C was as follows (12 cycles): denaturation at 98°C for 20 s; combined annealing and extension at 72°C (shuttle PCR) for 15 s, with a final extension at 72°C for 5 min. The products of the second PCR were combined (i.e., one pooled 2nd PCR product that included all samples), purified, excised and sequenced on the MiSeq platform using a MiSeq v2 Reagent Nano Kit for 2 × 150 bp PE. All sequences used in this study were deposited in DDBJ Sequence Read Archives (Submission ID = DRA005598; see Data Accessibility for further information).

### 2.5. Sequence read processing and taxonomic assignment

The overall quality of the MiSeq reads was evaluated, and the reads were assembled using the software FLASH with a minimum overlap of 10 bp [27]. The assembled reads were further filtered and cleaned, and the pre-processed reads were subjected to the clustering process and taxonomic assignments. The pre-processed reads from the above custom pipeline were dereplicated using UCLUST [28]. Those sequences represented by at least 10 identical reads were subjected to the downstream analyses, and the remaining under-represented sequences (with less than 10 identical reads) were subjected to pairwise alignment using UCLUST. If the latter sequences (observed from less than 10 reads) showed at least 99% identity with one of the former reads (i.e., no more than one or two nucleotide differences), they were operationally considered as identical (with the differences being attributed to sequencing or PCR errors and/or actual nucleotide variations in the populations).

The processed reads were subjected to local BLASTN searches against a custom-made database [29]. The custom-made database was generated as described in a previous study [14]. The top BLAST hit with a sequence identity of at least 97% and E-value threshold of 10^-5^ was applied for species assignments of each representative sequence. The detailed information about the above bioinformatics pipeline from data pre-processing through taxonomic assignment is available in the supplemental information in Miya et al. [3]. Also, an online version of this pipeline is available at http://mitofish.aori.u-tokyo.ac.jp/mifish.

### 2.6. Determination of the number of eDNA copies by quantitative PCR

The copy numbers of total fish eDNA were quantified using the SYBR-GREEN qPCR method using a StepOne-Plus™ Real-Time PCR system (Applied Biosystems, Foster City, CA, USA). SYBR-GREEN qPCR was conducted in a 10μL volume with a reaction solution that consisted of 5 μl of PowerUp™ SYBR^®^ GREEN Master Mix (Thermo Fisher Scientific, Wilmington, DE, USA), 0.6 μl of 5 μM MiFish-U-F/R primers (without adaptor), 10 μl of sterilized H_2_O, and 1.0 μl of DNA template. SYBR-GREEN qPCR was performed in triplicate for each eDNA sample, the standard dilution series, and PCR negative controls. The standard dilution series was prepared using DNA extracted from *Capoeta capoeta* (a freshwater fish species in Southeast Asia). We selected *C. capoeta* as the standard because the length of the MiFish region of this species is close to the average length in fish species (*C. capoeta* = 174 bp, the average length of the MiFish region = 173 bp). The thermal cycle profile after preconditioning for 2 min at 50°C and 2 min at 95°C was as follows (40 cycles): denaturation at 95°C for 3 s; annealing and extension combined at 60°C (shuttle PCR) for 30 s. Although MiFish primers predominantly amplify fish (e)DNA, we should note that the quantification by SYBR-GREEN qPCR may include non-fish eDNA because non-target sequences (e.g., sequences longer than the MiFish region) are sometimes amplified when using MiFish primers and because SYBR-GREEN qPCR does not distinguish between fish and non-fish eDNA. However, if the ratio of non-fish and fish amplicons does not drastically differ among samples, the SYBR-GREEN qPCR should reflect the dynamics (i.e., temporal fluctuation pattern) of total fish eDNA reasonably well. In all experiments, PCR negative controls showed no detectable amplification.

In addition, fish-species-specific eDNA was quantified by real-time TaqMan^®^ PCR according to Takahara et al. [8] using a StepOne-Plus™ Real-Time PCR system. The cytochrome *b* region of mitochondrial DNA was targeted for amplification from eDNA samples for each target species by using the following primer sets and associated probes, which were designed and confirmed to be able to amplify each target species-specifically [19 and Supplementary Information]. For the TaqMan qPCR analysis, Japanese anchovy (*Engraulis japonicus*) and Japanese jack mackerel (*Trachurus japonicas*) were chosen because they are abundant in the study area and standard dilution series were already available. For Japanese anchovy, primers Eja-CytB-Forward (5′-GAA AAA CCC ACC CCC TAC TCA-3′), Eja-CytB-Reverse (5′-GTG GCC AAG CAT AGT CCT AAA AG-3′), and Eja-CytB-Probe (5′-FAM-CGC AGT AGT AGA CCT CCC AGC ACC ATC C-TAMRA-3′) were used. For Japanese jack mackerel, primers Tja-CytB-Forward (5′-CAG ATA TCG CAA CCG CCT TT-3′), Tja-CytB-Reverse (5′-CCG ATG TGA AGG TAA ATG CAA A-3′), and Tja-CytB-Probe (5′-FAM-TAT GCA CGC CAA CGG CGC CT-TAMRA-3) were used. The length of the PCR amplicon produced using the primer set was 115 bp and 127 bp for Japanese anchovy and for Japanese jack mackerel, respectively. PCR was conducted in a 15μl volume containing each primer at 900 nM, TaqMan^®^ probe at 125 nM, and 2 μl of sample DNA in 1 × PCR master mix (TaqMan^®^ gene expression master mix; Life Technologies, Carlsbad, CA, USA). A dilution series of standards was prepared for quantification and analyzed at the concentration of 3 × 10^1^ to 3 × 10^4^ copies per well in each experiment to obtain standard curves. The standards were pTAKN-2 plasmids containing commercially synthesized artificial DNA that had the same sequence as the amplification region of each species. The thermal cycle profile after preconditioning for 2 min at 50°C and 10 min at 95°C was as follows (55 cycles): denaturation at 95°C for 15 s; combined annealing and extension at 60°C (shuttle PCR) for 60 s. qPCR was performed in triplicate for each eDNA sample, standard dilution series, and PCR negative controls. In all experiments, PCR negative controls showed no detectable amplification.

### 2.7. Statistical analyses

For all analyses, the free statistical environment R was used [30]. Our statistical analyses consisted of three parts: (1) linear regression analysis to examine the relationship between sequence reads and the copy numbers of the standard DNA for each sample, (2) the conversion of sequence reads of non-standard fish eDNA to calculated copy numbers using the result of the linear regression for each sample, and (3) the comparison of eDNA copy numbers quantified by MiSeq and qPCR.

Linear regressions were performed using the *lm* function in R, and used to examine how many sequence reads were generated from one (e)DNA copy through the library preparation process for MiSeq. Note that a linear regression between sequence reads and standard DNAs was performed for each sample and the intercept was set as zero. The regression equation was: MiSeq sequence reads = regression slope × the number of standard DNA copies [/μl]. The number of linear regressions performed was 52 (= the number of eDNA samples), and thus 52 regression slopes were estimated in total (see Fig. S1).

The sequence reads of non-standard fish eDNAs were converted to calculated copy numbers using a sample-specific regression slope estimated by the first analysis. The number of non-standard eDNA copies was estimated by dividing MiSeq sequence reads by a sample-specific regression slope (i.e., the number of DNA copies = MiSeq sequence reads/regression slope; hereafter, this equation is referred to as ‘correction equation’). The estimated numbers of non-standard fish eDNA copies are hereafter referred to as ‘calculated copy numbers’, and this method itself (i.e., from an inclusion of standard DNA to the conversion of sequence reads using a correction equation) is hereafter referred to as ‘qMiSeq’.

Calculated copy numbers by qMiSeq were compared with copy numbers estimated by qPCR (see above sections for detailed qPCR method) by using four approaches. First, raw values (non-standardized copy numbers) were compared using linear regressions (i.e., first approach). Because there were significant outliers, linear regressions were again performed by excluding the outliers (i.e., second approach). In addition, because the distribution of calculated copy numbers was highly right-skewed (i.e., many samples with low copy numbers and few samples with high copy numbers), log-transformation (base = 2) was applied after adding 0.5 to the raw values. The log-transformed values were further compared using linear regressions (i.e., third approach). Lastly, a Bland-Altman plot (difference plot) [31] was constructed (for raw and log-transformed values) to measure the agreement between qMiSeq and qPCR using BlandAltmanLeh package [32] (i.e., fourth approach). Linear relationships were considered significant if *P* values were smaller than 0.05.

All R codes, original data tables used for the analyses, and figure generation codes are available in https://github.com/ong8181/eDNA-qmiseq.

## 3. Results and discussion

### 3.1. Relationship between the copy numbers and sequence reads of the standard DNA

The sequence reads of the internal standard DNAs were significantly positively correlated with the copy numbers of those DNAs (Fig. 2a, b, Fig. S1). A regression line was drawn for each eDNA sample, and therefore the number of regression lines equaled the number of eDNA samples (= 52; Fig. S1). R^2^ values of the regression lines ranged from 0.71 to 0.98, and more than 80% of regression lines showed R^2^ values higher than 0.9 (Fig. 2c; see Fig. S2 for regression residuals), suggesting that the number of sequence reads was proportional to the number of DNA copies in a single sample and that the slopes of the regression lines (i.e., sequence reads per DNA copy) can be used to convert sequence reads to the numbers of DNA copies. Interestingly, the slopes of the regression lines were highly variable, ranging from 0 to 54.1 (which corresponded to eDNA samples collected on 2015/6/16 and 2015/4/21, respectively), with a median value of 24.6 (Fig. 2d, Fig. S1). Low slope values (e.g., 0, or close to 0) indicate that internal standard DNAs were not efficiently amplified even if the number of DNA copies added was large, suggesting the presence of PCR inhibitor(s) (e.g., humic substance) in the eDNA samples. Also, these variations of the slope suggested that the degree of PCR inhibition varies depending on the eDNA sample.

**Figure 2.**
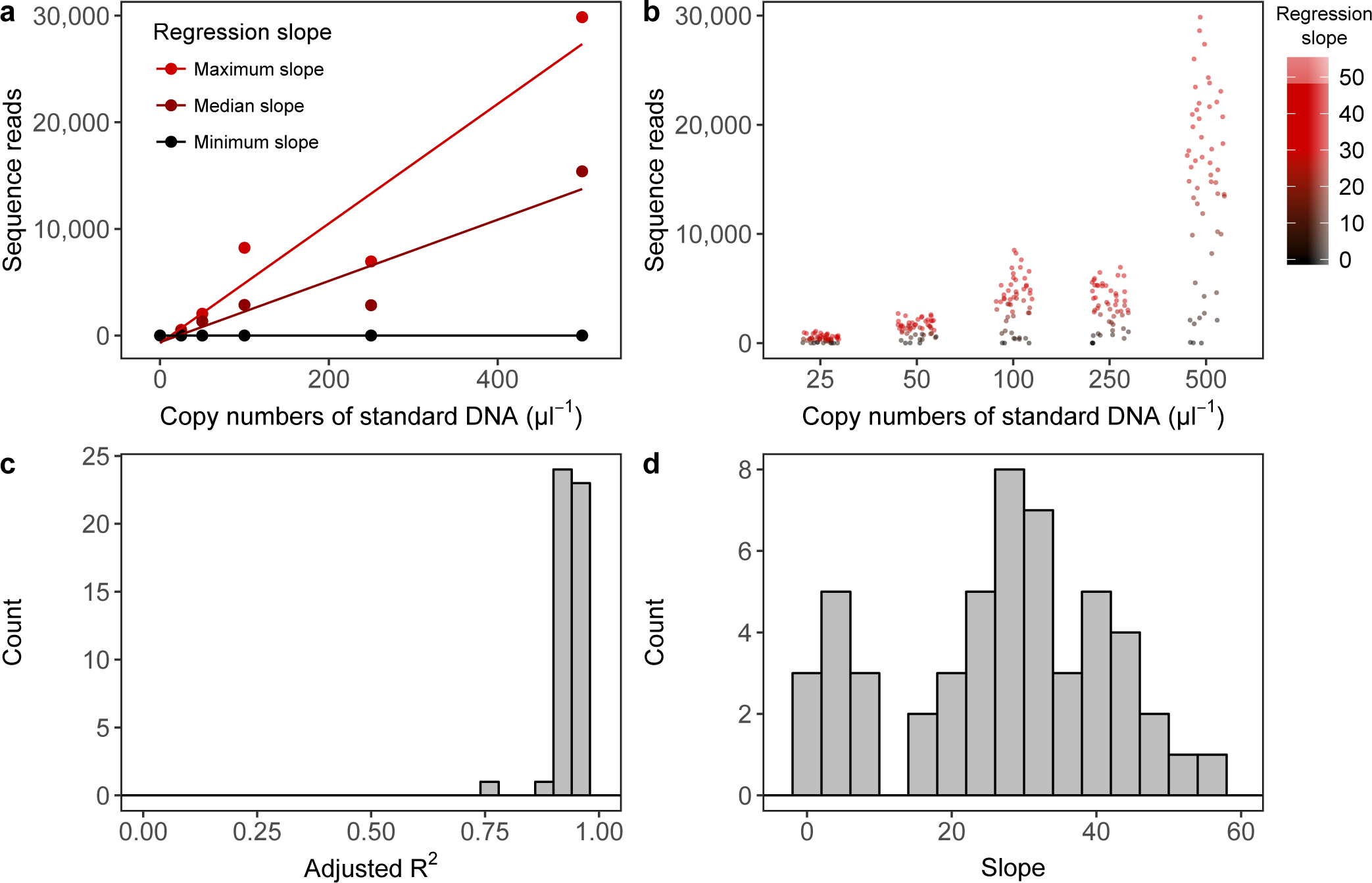
Summary of regression lines constructed using the number of copies added and sequence reads of internal standard DNAs. Examples of a regression line (**a**). Regression lines with the maximum, median, and minimum slopes are indicated as examples of the relationships. The line indicates the regression line between the copy numbers of standard DNA (copiesμl) and sequence reads. The intercept of the regression line was set as zero. Distributions of sequence reads of internal standard DNAs (**b**). The intensity of red colour indicats the slope of regression line. Distribution of adjusted R^2^ of the regression line (**c**). Note that a regression line was drawn for each eDNA sample and that the number of standard curves is equal to the number of eDNA samples (*N* = 52). Distribusion of slopes of regression lines (**d**).

### 3.2. Quantification of the copy number using sequence reads and correction equations, and comparison of the calculated copy number with the copy number quantified by qPCR

MiSeq sequence reads of each sample were converted using each correction equation (i.e., the number of eDNA copies [copiesμl] = MiSeq sequence reads / a sample-specific regression slope; the copy numbers and this method itself are referred to as ‘calculated DNA copies [copiesμl]’, and ‘qMiSeq’, respectively). Then, calculated DNA copies were compared with the number of DNA copies quantified by qPCR (Fig. 3 [all regression lines were significant, *P* < 0.05] and Table S2 and S3). The numbers of eDNA copies estimated by qMiSeq and qPCR were significantly and positively correlated with each other for total fish eDNA (all data included, Fig. 3a; outliers excluded, Fig. 3b). For the total fish eDNA, the number of eDNA copies quantified by qPCR (mean copy number = 683 copiesμ/l) was higher than that quantified by qMiSeq (mean copy number = 139 copiesμ/l). This is not surprising because we excised target amplicon fragments (about 370 bp, including MiSeq adaptor), and we discarded non-target amplified fragments (e.g., longer and unknown amplicons), which were included in the quantification by the SYBR-GREEN assay, before MiSeq sequencing. Regarding the eDNA of Japanese anchovy and Japanese jack mackerel, we found that the number of eDNA copies quantified by qMiSeq was similar to that obtained by qPCR (i.e., regression lines were close to the 1:1 line in Fig. 3c–f, regardless of the inclusion/exclusion of outliers; for the relationships between sequence reads and copy numbers quantified by qPCR, see Fig. S3). In addition, the Bland-Altman plot for the raw values also showed that the differences in the copy numbers quantified by the two methods were not significantly different from zero (Fig. S4). These results suggested that eDNA metabarcoding with the inclusion of internal standard DNAs reasonably quantified the number of eDNA copies.

**Figure 3.**
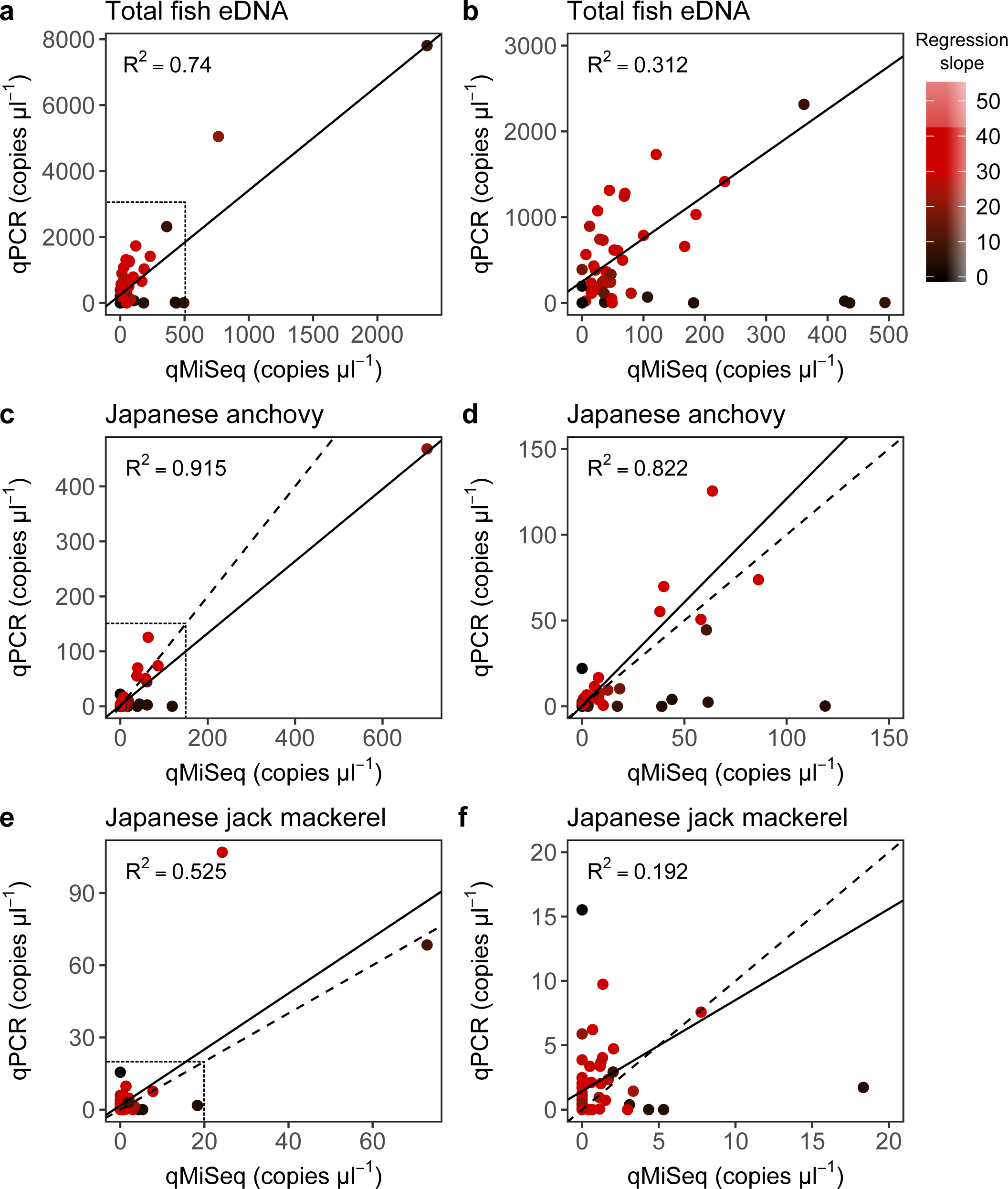
Relationship between the number of eDNA copies quantified by qPCR and that by qMiSeq. Correlations for the total fish eDNA (all data, **a**; enlarged figure, **b**), Japanese anchovy (all data, **c**; enlarged figure, **d**) and Japanese jack mackerel (all data, **e**; enlarged figure, **f**). Dashed and solid lines indicate 1:1 line and linear regression line, respectively. Regression lines in the enlarged figures were drawn by excluding outliers. All regression lines were significant (*P* < 0.05). Dotted boxed regions in **a**, **c**, and **e** correspond to the range of the graphs in **b**, **d**, and **f**, respectively. The intensity of red colour indicates the slope of the regression line (=correction equation) used to convert sequence reads to the calculated copy numbers.

Although the calculated DNA copies generally corresponded well with the eDNA copy numbers estimated by qPCR, the calculated DNA copies of some samples were much higher than the copy numbers obtained by qPCR (i.e., for the points close to the x-axis in Fig. 3). These samples showed relatively small values of the slopes of regression lines between the sequence reads and quantity of the standard DNAs (i.e., corresponded to points with darker colour in Fig. 3), suggesting that there was inhibition of PCR in these samples. qMiSeq can control for PCR-inhibition effects in the estimation of eDNA copy because correction equations already take the influence of PCR inhibition into account, which may be an advantage of this method compared with qPCR. Conversely, we would suggest that qPCR could not reliably quantify the number of eDNA copies when the influence of PCR inhibitors was strong. Theoretically, influences of PCR inhibition could also be tested by (multiplex) qPCR [e.g., 33,34], but multiplex qPCR may require some additional experimental procedures, and thus could be more time-consuming and costly.

Some samples showed much lower eDNA copy numbers of Japanese jack mackerel when quantified by qMiSeq than when quantified by qPCR (i.e., points close to the y-axis; Fig. 3f). This inconsistency might have been due to the low eDNA copy number of Japanese jack mackerel (all samples showed less than 100 copiesμl, and most samples showed less than 10 copiesμl). qMiSeq might not be able to quantify such low numbers of eDNA copies accurately because the lowest copy number of internal standard DNA added was 25 copies/μl If the copy number of internal standard DNA had been much lower, more accurate quantification would have been achieved by qMiSeq. Furthermore, the difference in number of PCR cycles between qPCR (40-55 cycles) and the 1st PCR of MiSeq library preparation (35 cycles) might contribute to the different sensitivities (i.e., detection limits) of these methods.

Although the above analyses suggested that there is a good agreement between the two methods, the distributions of calculated copy numbers as well as copy numbers estimated by qPCR were right-skewed (i.e., many low copy numbers and few high copy numbers), and thus we further compared the copy numbers after log-transformation of the raw values. Samples with regression slopes lower than 10 were excluded from this analysis because they suggested that there had been significant PCR inhibition during the qPCR measurements, as discussed above. We found that there were positive and significant linear relationships between the copy numbers quantified by qMiSeq and qPCR even after the log-transformation (*P* < 0.05; Fig. 4a-c). Bland-Altman plots also suggested that there is a good agreement between the two methods (i.e., 95% confidence intervals include zero; Fig. 4e, f). Taken together, these results suggested that eDNA metabarcoding with internal standard DNA enabled simultaneous quantification and identification of fish eDNA. The appropriate range of the copy numbers of the internal standard DNAs should, however, be carefully determined depending on the range of the target eDNA copy numbers in environmental samples.

**Figure 4.**
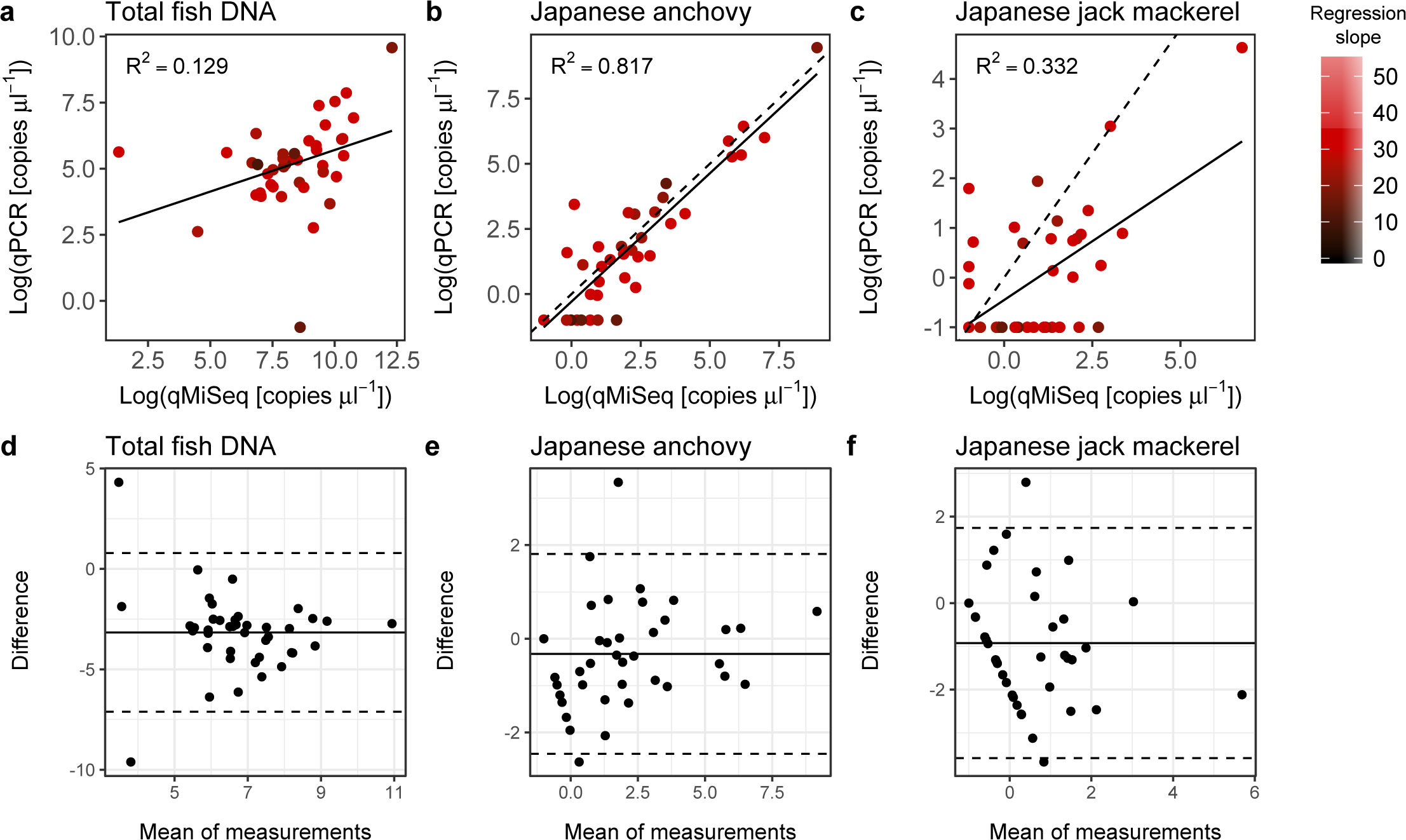
Relationship between log-transformed copy numbers quantified by qPCR and qMiSeq for the total fish eDNA (**a**), Japanese anchovy (**b**), and Japanese jack mackerel (**c**). A solid line is a linear regression line (all lines are significant; *P* < 0.05). The intensity of red colour indicates the slope of the regression line (i.e., correction equation) used to convert sequence reads to the copy numbers. Bland-Altman plot for log-transformed copy numbers of the total fish eDNA (**d**), Japanese anchovy (**e**), and Japanese jack mackerel (**f**). Dashed lines indicates 95% upper and lower limits, and solid line indicates mean values. Note that, although it is not significant, the Bland-Altman plot for the log-transformed copy numbers of the total fish eDNA (**d**) showed that the calculated copy numbers by qMiSe tends to be smaller than those by qPCR probably because of the removal of non-target amplified fragments before MiSeq sequencing (see discussion in the main text).

### 3.3. Temporal dynamics of Maizuru-bay fish eDNA revealed by eDNA metabarcoding

The temporal dynamics of the fish eDNA quantified by qMiSeq generally corresponded well with those quantified by qPCR (Fig. 5). For the total fish eDNA, the highest eDNA concentrations were found on 25th August and 24th November by qPCR, and peaks were also detected on those dates by qMiSeq (Fig. 5a). For Japanese anchovy eDNA, the highest eDNA concentration was found on 24th November by qPCR, and the peak was also detected on this date by qMiSeq (Fig. 5b). For Japanese jack mackerel eDNA, one of the highest eDNA concentrations (on 25th August) found by qPCR was also found by qMiSeq. However, another peak found by qPCR (on 23rd February) was not detected by qMiSeq (Fig. 5c), probably due to the above-mentioned technical issues in eDNA metabarcoding with internal standard DNA.

**Figure 5.**
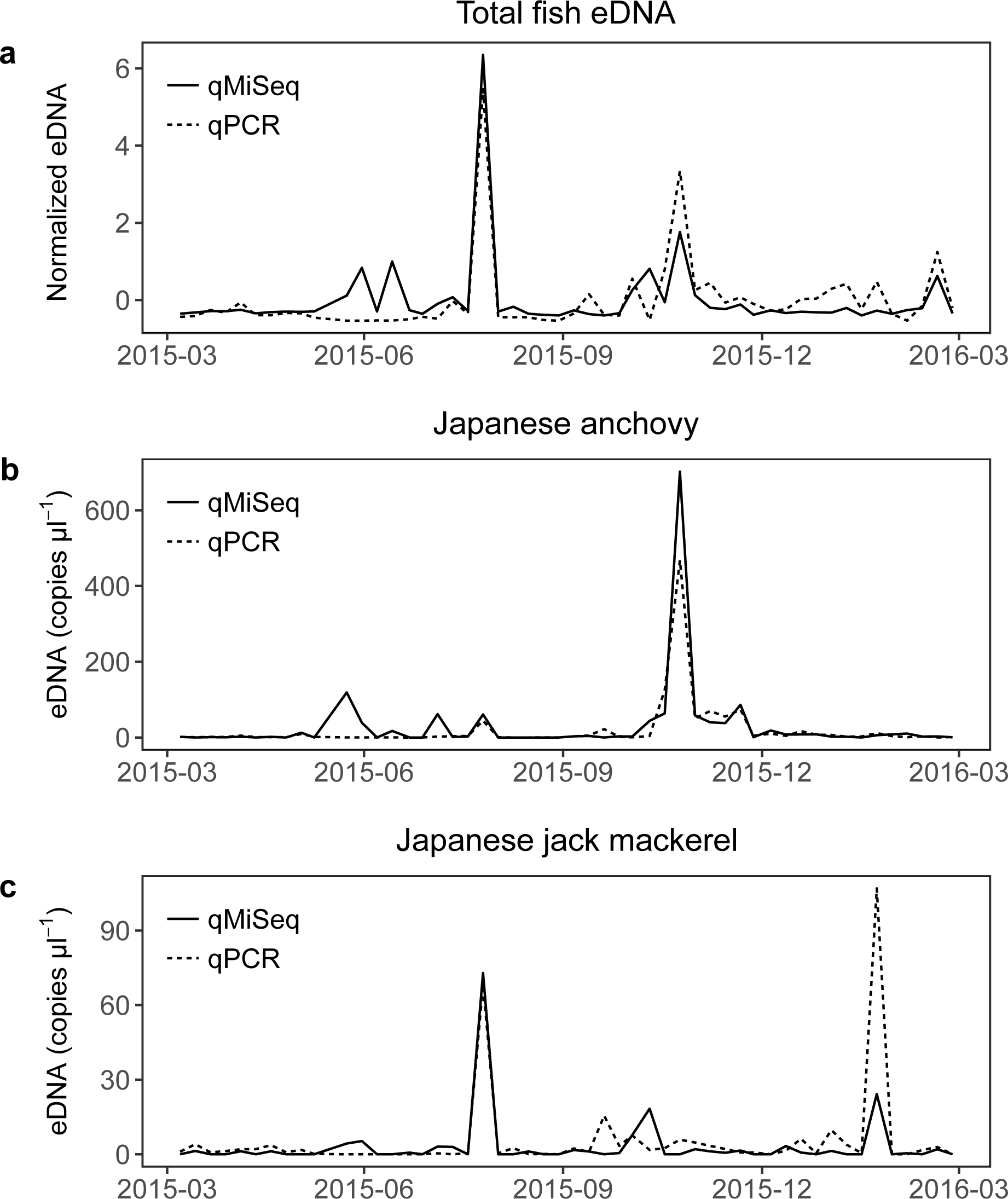
Dynamics of the total fish eDNA (**a**), Japanese anchovy (**b**) and Japanese jack mackerel (**c**) quantified by eDNA metabarcoding and qPCR. Solid and dashed lines indicate the number of eDNA copies quantified by eDNA metabarcoding and qPCR, respectively. Note that the copy numbers of total fish eDNA were normalized to have zero and unit variance.

The results obtained in the present study suggest that qMiSeq can reasonably recover the dynamics of fish eDNA. Because eDNA metabarcoding can detect many species (sometimes more than 100 species) in a single run [3], this method enables simultaneous quantifications of eDNA derived from many fish species. In the present study, we detected more than 70 fish species from 52 eDNA samples collected from April 2015 to March 2016 in Maizuru Bay, Kyoto, Japan (Supplementary text, Table S2), which is generally consistent with long-term direct visual observations, e.g., fortnightly-performed visual census over 5 years detected a total of 83 fish species [1,26].

### 3.4 Quantitative and multispecies fish eDNA monitoring

Our method enables the generation of a quantitative time series of eDNA of these fish species by a single MiSeq run, and as an example, an eDNA time series of the 10 most abundant fish species in terms of eDNA concentration is shown in Fig. 6. Because eDNA copy numbers may be a rough index for fish biomass/abundance [11], such a multispecies quantitative time series, which can readily be obtained if the qMiSeq method is used, may provide valuable information about the dynamics of fish populations in the sampling area. Indeed, the eDNA time series measured here by qMiSeq were ecologically interpretable, suggesting that eDNA monitoring using our method would provide ecologically meaningful information on the dynamics of a natural fish community, at least in our case (see Supplementary Information and Table S4).

**Figure 6.**
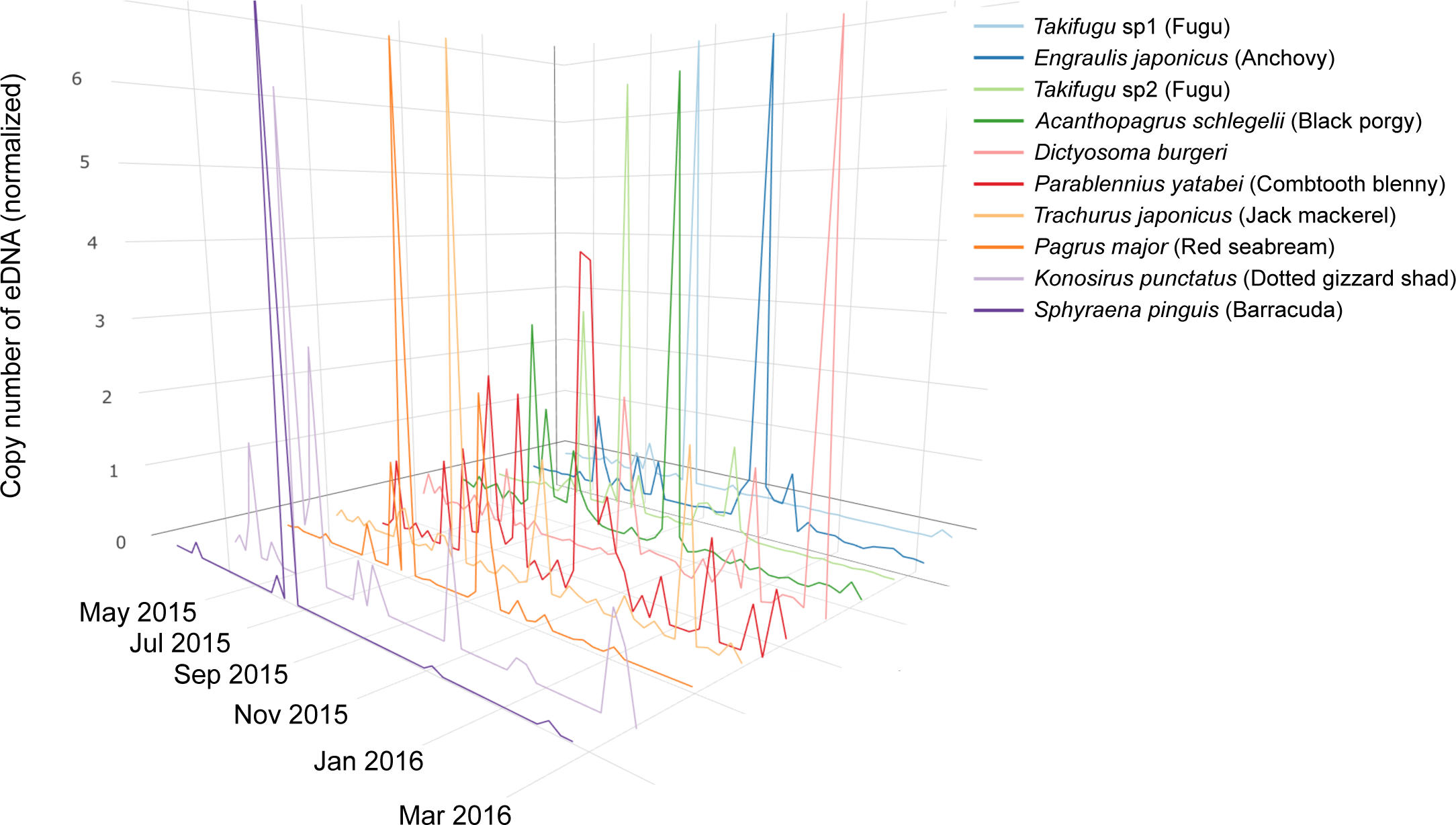
Quantitative and multispeies fish eDNA time series in Maizuru Bay, Kyoto, Japan. Time series of eDNA of 10 dominant fish species. In the eDNA analysis, two *Takifugu* species were detected as dominant species, and were designated *Takifugu* sp1 and sp2. Representative sequence of *Takfugu* sp1 is highly similar to that of *T niphobles/T snyderi* (>99% identity). Representative sequene of *Takifugu* sp2 is identical with that of *T pardadalisiT xanthopterusiT poecilonotus* (100% identity). Different colours indicate different fish species. The number of eDNA copies were normalized to have zero mean and unit variance.

As eDNA metabarcoding has been recognized as an efficient approach in species detection and biodiversity assessment [5,12–15,35], its use as a biodiversity monitoring tool has been increasing [12,17,36]. The authors of those monitoring studies performed periodic water samplings and generated eDNA time series, and showed that temporal fluctuations in species (or OTU) richness and detection probability of eDNA of a target taxa [12,17,36,37] were in good agreement with temporal fluctuations in other reliable data (e.g., visual census). However, because of a lack of a quantitative method of evaluating eDNA metabarcoding, only qualitative information about eDNA (e.g., presence/absence, rank of eDNA sequence reads, and species/OTU diversity) has been reported for the comparisons between eDNA monitoring data and other monitoring data. The use of sequence reads as a quantitative index of the abundance/biomass of target organisms may partly solve this problem [20]. However, the number of sequence reads per sample (or per species) may change dramatically depending on experimental conditions such as the number of samples multiplexed, final library concentrations and sequence reagents, and thus rigorous comparisons between samples originated from different experiments/studies were difficult.

The quantities of internal standards are precisely known, and thus the use of an internal standard would enable rigorous and quantitative comparisons even between different experiments/studies, which would facilitate the use of eDNA metabarcoding as a tool for biodiversity monitoring [21,25 and this study]. Furthermore, our method, i.e., the addition of purified DNA fragments, would be less time- and effort-consuming than the use of tissues of standard organisms as internal standards [21,25] because the preparation of standard organisms/tissues is sometimes difficult. In future studies, the use of artificial fish sequences, that are not identical to the sequences of any other fish species in the world, should be considered because it would be applicable to any water sample and would further increase the efficiency of our method.

## 4. Conclusion

In the present study, we showed that eDNA metabarcoding performed with the inclusion of internal standard DNA enables simultaneous determination of the quantity and identity of eDNA derived from multiple fish species. Because the traditional species-specific qPCR allows quantification of eDNA from only one fish species in a single experiment, our method is much more efficient compared with qPCR. In addition, our method can take effects of PCR inhibition into account. Although it should be mentioned that fish eDNA copy numbers are still only a rough index of fish biomass/abundance (or population size) and this problem should be addressed in a future study, our results show that eDNA metabarcoding with the inclusion of internal standard DNAs can be a promising tool to monitor fish biodiversity. Our method will improve the efficiency of obtaining data, and may contribute to more effective resource management and ecosystem monitoring.

## Ethics

The experiments were conducted in accordance with the guidelines of Regulation of Animal Experimentation at Kyoto University. Water sampling permission in or around the pier was not needed.

## Data accessibility

Sequences are deposited in DDBJ Sequence Read Archive (DRA): Accession numbers are: DRA005598 (Submission ID), PRJDB5570 (BioProject ID) and SAMD00075651-SAMD00075720 (BioSample ID). All R codes and original data table used for the analyses are available in GitHub (https://github.com/ong8181/eDNA-qmiseq).

## Authors’ contributions

MU conceived and designed research; HM and RM performed sampling; MU, HM, RM, TS, MM, SS, HY and TM performed experiments; MU analyzed data; MU wrote the early draft and completed it with significant inputs from all authors.

## Acknowledgements

We would like to thank Aina Tanimoto for help with the collection and filtration of water samples.

